# Enhanced release of ciliary extracellular vesicles suppresses cell migration and promotes cell aggregation

**DOI:** 10.1101/2025.11.30.691462

**Authors:** Tetsuhiro Hara, Ryota Nakazato, Kenji Shirakawa, Faryal Ijaz, Kenichiro Uemura, Shinya Takahashi, Koji Ikegami

## Abstract

A primary cilium is a hair-like organelle that protrudes from the cell surface in many cell types. Growing evidence indicates that extracellular vesicles are released from primary cilia, and research is increasingly focused on defining the functions of these cilia-derived extracellular vesicles (EVs). EVs are known to modulate the behavior of various cancer cells, and structural and functional abnormalities in primary cilia have been reported in multiple cancer types. We previously demonstrated that PANC-1 cells, a human pancreatic ductal adenocarcinoma cell line, acquire enhanced primary cilia formation after surviving solitary culture conditions, and that their cilia contribute to tumor-like cell mass formation. Here, we explored part of the underlying mechanism of this phenotype by investigating the contribution of EVs released from the primary cilia of PANC-1 cells. PANC-1 clones generated by limiting dilution exhibited enhanced ciliogenesis and distinct ciliary morphologies compared with parental cells. These clones also released higher levels of cilia-derived EVs, including an expanded population of freely floating EVs within the culture environment. Biochemical analyses further showed that this increase was selective for primary cilia-derived EVs rather than reflecting a global rise in total EV production. Functionally, EV fractions enriched in cilia-derived EVs suppressed parental PANC-1 cell migration, altered cell morphology, and promoted cell aggregation, mimicking key behavioral traits of solitary condition-surviving PANC-1 clones. Together, these findings identify enhanced release of primary cilia-derived EVs as a distinct feature of PANC-1 cells adapted to solitary growth and suggest their potential involvement in the malignant and metastatic behaviors of pancreatic cancer.

## Introduction

A primary cilium is a microtubule-based sensory organelle that is formed on the surface of almost all vertebrate cell types and functions to sense mechanical and chemical stimuli (Garcia-Gonzalo and Reiter, 2012). The dynamics of primary cilia is crucial for the regulation of cellular homeostasis including cell cycle control (Plotnikova *et al*., 2009). Proper ciliary function is essential for accurate cell and tissue differentiation, as well as for normal organ and organismal development (Corbit *et al*., 2005; Nusse, 2003; Pazour and Witman, 2003; Singla and Reiter, 2006). A group of disorders collectively termed ciliopathies, which involve structural defects or dysfunction of primary cilia, affect the architecture and function of multiple organs (Badano *et al*., 2006).

Recent studies have demonstrated that the tips of primary cilia in cultured mammalian cells can be severed and released into the extracellular space as extracellular vesicles (Nager *et al*., 2017; Phua *et al*., 2017; Wang *et al*., 2019). Although primary cilia have long been recognized as sensors of extracellular stimuli, such as hormones, growth factors, and fluid flow (Anvarian *et al*., 2019), the physiological roles of primary cilia-derived vesicles in mammalian cells remain largely unclear (Ikegami and Ijaz, 2021). In contrast, similar EVs released from cilia or flagella of model organisms such as *Chlamydomonus* and *C. elegans* have been extensively characterized (Wang *et al*., 2014; Wang *et al*., 2024; Wood *et al*., 2013). Notably, one study has reported that a type of glioma cell exhibits enhanced proliferative potential when cultured in medium enriched with primary cilia-derived EVs (Hoang-Minh *et al*., 2018), suggesting that such vesicles may mediate intercellular communication and could be associated with the poor prognosis observed in ciliated tumor cells.

Primary cilia have also been implicated in carcinogenesis. In many cancer types, including pancreatic ductal adenocarcinoma (Chao *et al*., 2022), primary cilia are frequently lost (Schimmack *et al*., 2016). This loss is thought to occur during the early stages of oncogenic transformation and tumorigenesis (Menzl *et al*., 2014; Seeley *et al*., 2009). Inhibition of ciliogenesis has been reported to promote the malignant transformation of normal pancreatic ductal epithelial cells (Deng *et al*., 2018; Kobayashi *et al*., 2020). The loss of primary cilia is therefore considered to be strongly correlated with cancer progression (Hassounah *et al*., 2012; Paul *et al*., 2022; Seeger-Nukpezah *et al*., 2013). However, a clinical study of PDAC patients reported that the presence of primary cilia is associated with an increased rate of lymph node metastasis and serves as an independent poor prognostic factor (Emoto *et al*., 2014). Interestingly, similar context-dependent and even contradictory roles of primary cilia have been reported in other cancer types as well (Han *et al*., 2009; Wang *et al*., 2021). In recent years, an increasing number of studies have suggested a relationship between primary cilia and resistance to chemotherapy (Chao *et al*., 2022; Jenks *et al*., 2018; Khan *et al*., 2018; Kim *et al*., 2022; Lee, 2023).

PANC-1 is a cell line originally established from a primary PDAC tumor (Lieber *et al*., 1975). We previously reported that the ability to form primary cilia is enhanced in PANC-1 cells cultured under solitary conditions. Furthermore, we demonstrated that primary cilia promote the formation of tumor-like cell clusters in PANC-1 cells (Shirakawa *et al*., 2025). In this study, we explored the underlying mechanisms by which primary cilia promote cell cluster formation in PANC-1 cells cultured under the solitary condition. We found that solitary-cultured PANC-1 cells not only exhibit increased ciliogenesis but also release larger amounts of primary cilia-derived EVs. Moreover, we demonstrated that these EVs suppress PANC-1 cell migration while altering cell morphology and promoting the formation of cell aggregates.

## Materials and methods

### Cell culture

A human pancreatic cancer cell line, PANC-1, was obtained from the RIKEN Bioresources Cell Bank (Ibaraki, Japan). Cells were maintained in RPMI-1640 (Wako) supplemented with 10% fetal bovine serum (FBS; Gibco) and cultured at 37 °C in a humidified incubator with 5% CO_2_. For routine passaging, cells were seeded onto 100-mm culture dishes and harvested using 0.25% trypsin-EDTA. Cell numbers were determined with a hemocytometer. Phase-contrast images of cultured cells were acquired using an inverted microscope (AE3000, SHIMADZU RIKA).

### Antibodies

The following primary antibodies were used for immunofluorescence (IF) and/or Western blotting (WB): ARL13B (rabbit polyclonal, 1:1,000 for IF and WB; 17711-1-AP, Proteintech), ARL13B (mouse monoclonal, 1:50 for IF; 66739-1-Ig, Proteintech), acetylated tubulin (mouse monoclonal 6-11B-1, 1:50,000 for IF; T7451, Sigma), FOP (mouse monoclonal, 1:10,000 for IF; H00011116-M01, Abnova), ALIX (rabbit polyclonal, 1:1500 for WB; 12422-1-AP, Proteintech), GAPDH (mouse monoclonal, 1:3,000 for WB; MAB374, Millipore), KIF3A (rabbit polyclonal, 1:1000 for WB; 13930-1-AP, Proteintech), IFT88 (rabbit polyclonal, 1:1,500 for WB; 13967-1-AP, Proteintech). Secondary antibodies include goat anti-mouse IgG (H+L) highly cross-adsorbed secondary antibody, Alexa Fluor 488 (1:1,000; A-11029; Thermo Fisher Scientific), goat anti-rabbit IgG (H+L) highly cross-adsorbed secondary antibody, Alexa Fluor 568 (1:1,000; A-11036; Thermo Fisher Scientific) for IF, and horseradish peroxidase-conjugated antibodies for WB (1:10,000; 111-035-003; Jackson Immuno Research Laboratories). To minimize appearance of false particles, all primary and secondary antibodies used for EVs detection were centrifuged at 20,000 × g for 30 min prior to use to remove protein or fluorophore aggregates.

### Immunofluorescence staining

Cells grown on coverslips (C018001; MATSUNAMI) were fixed with 4% paraformaldehyde (PFA, pH 7.5; 162-16065; Wako) for 20 min at room temperature. Following fixation, cells were blocked and permeabilized with PBS containing 5% normal goat serum and 0.1% Triton X-100 for 1 h at room temperature. Cells were then incubated overnight at 4 °C with primary antibodies diluted in the same blocking buffer. After washing with PBS, cells were incubated with Alexa Fluor-conjugated secondary antibodies and DAPI (1 µg/ml; DOJINDO) for 1 h at room temperature. Samples were mounted on glass slides using VECTASHIELD mounting medium (Vector Laboratories, Burlingame, CA). Fluorescence images were acquired using an epifluorescence microscope (DMI3000B, Leica) equipped with 100× objective lens, N.A. 1.40 (11506378; Leica) or a laser scanning confocal microscope (STELLARIS 5, Leica) equipped with 63× objective lens, N.A. 1.40 (11506350; Leica). Quantitative image analysis was performed using ImageJ software (National Institutes of Health, Bethesda, MD, USA) (Schindelin *et al*., 2012; Schneider *et al*., 2012).

### Collection of primary cilia-derived EVs

The collection of primary cilia-derived EVs in this study was performed based on the standard method for collecting EVs by ultracentrifugation (Théry *et al*., 2006). FBS used for EVs collection was pre-depleted by ultracentrifugation at 100,000 × g to remove FBS-derived small EVs such as exosomes and ectosomes. Forty-eight hours after seeding, the culture medium was replaced with medium containing 1% pre-cleared FBS. Conditioned culture medium was collected 20-24 h later. EVs released into the culture medium were collected using a three-step differential centrifugation protocol. First, culture medium was centrifuged at 2,000 × g for 20 min to remove dead cells. The resulting supernatant was centrifuged at 10,000 × g for 30 min to remove debris as pellet. The resulting supernatant was ultracentrifuged at 100,000 × g for 3 h 10 min using an MLA-55 angle rotor (Beckman). Pellets were rinsed once with PBS and subsequently used for Western blotting. For collecting EVs used in wound-healing assay and actin staining, ultracentrifugation was performed using MLS-50 swing rotor (Beckman) with a 250 µl glycerol cushion placed at the bottom of the tubes to prevent deformation and rupture of EVs. Following ultracentrifugation, supernatant was carefully removed, and the EV pellets were resuspended in RPMI-1640 medium containing 1% pre-cleared FBS for use in each experiment. The EVs amount was standardized by the initial number of cells seeded for the collection of conditioned medium, as the proliferation rates of clone #70 and #76 are comparable to those of parental PANC-1 cells (Shirakawa *et al*., 2025). Parental PANC-1 cells and clone #70 and #76 were seeded at a density of 10.6 × 10^4^ cells/cm^2^.

### Western blotting analysis

Cells and EV pellets were lysed with 1× sodium dodecyl sulfate-polyacrylamide gel electrophoresis (SDS–PAGE) sample buffer (Wako) and heated at 95 °C for 5 min. Samples were loaded onto an acrylamide gel to keep the ratio of EV to cell lysate identical for all proteins examined. Proteins separated in SDS-PAGE were then transferred onto polyvinylidene difluoride (PVDF) membranes (Merk Millipore, Burlington, MA, USA). Membranes were blocked with 5% bovine serum albumin (BSA) prepared in Tris-buffered saline containing 0.1% Tween-20 (TBST) at room temperature for 1 h, followed by incubation with primary antibodies diluted with 1% BSA/TBST at 4 °C overnight. Following washes with TBST, membranes were incubated with horseradish peroxidase-conjugated secondary antibodies at room temperature for 1 h. Protein bands were visualized using ECL prime (GE Healthcare, Chicago, IL, USA) and detected with VersaDoc imaging system (Bio-Rad, Hercules, CA). Band intensities in Western blotting were measured using the “Gels” tool in ImageJ. All experiments were independently performed at least three times to ensure reproducibility.

### Knockdown with RNAi

The Dicer-substrate small interfering RNA (DsiRNA) duplexes used were previously reported (Shirakawa *et al*., 2025). Clone #70 were seeded into 35-mm or 100-mm culture dishes at a density of 56,000-57,000 cells/cm^2^. Cells were transfected with 1 nM DsiRNA and 2 nM DsiRNA on day 1 and day 3 using Lipofectamine RNAiMAX (Invitrogen) with the manufacturer’s instruction. On day 5, the culture medium was replaced with medium containing 1% pre-cleared FBS. On day 6, cells were either fixed or solubilized for subsequent immunofluorescence or Western blotting analysis. The conditioned medium was collected to precipitate EVs for Western blotting.

### Wound healing assay

Confluent parental PANC-1 cells were scratched using a sterile 200 µl plastic pipette tip, and the culture medium was immediately replaced with medium containing EVs collected from conditioned media of either parental PANC-1 cells or clone #70 and #76. Images of the wound area were captured at 0 and 48 h post-scratch using a phase-contrast microscope. Wound closure was quantitated using ImageJ software.

### Actin staining

Parental PANC-1 cells were treated with culture medium containing EVs collected from condition media of either parental PANC-1 cells or clone #70 and #76. After 48 h of treatment, cells were fixed with 4% PFA. The cells were then incubated for 2 h at room temperature with PBS containing AlexaFluor 488-conjugated phalloidin (1:1,000; A12379, Invitrogen) and DAPI (1:1,000) to stain F-actin and nuclei, respectively. Fluorescence images were acquired using STELLARIS 5 confocal microscope. For quantification, five random fields per sample were captured, and the proportion of round-shaped cells was calculated relative to the total number of DAPI-stained cells.

### Image analyses

Quantitative analyses of microscopy images were performed using ImageJ software. Cells bearing primary cilia were determined by counting DAPI-stained nuclei co-localized with ARL13B-positive cilia. Primary cilia were defined as ARL13B-positive structures anchored by FOP-positive centrioles. The morphology of primary cilia was classified according to the following criteria: ‘Bulged’, primary cilia with tip diameters of 1 µm or greater; ‘Emission’, presence of ARL13B-positive particles within 2 µm of the primary cilia tip; ‘Very short’, primary cilia with lengths ranging from 0.5 to 1 µm; ‘Branched’: two primary cilia formed from a single centriole.

Acetylated tubulin (Ac-tubulin) positivity was determined by classifying bulged tips with fluorescence intensities above a defined threshold. The threshold for Ac-tubulin positivity was set as the 99th percentile of the nuclear Ac-tubulin signal intensity, based on the assumption that Ac-tubulin is absent from the nucleus. Fluorescence intensities of nuclear Ac-tubulin were measured using the ‘Analyze Particles’ tool of ImageJ.

Primary cilia-derived EVs were quantified by counting ARL13B-positive, FOP-negative particles. Images were acquired using either an epifluorescence microscope (DMI3000B, Leica) or a confocal microscope (FV1000, Olympus). EV localization on cells was analyzed using the “Orthogonal Views” tool ImageJ. Primary cilia-derived EVs were defined and quantified as follows: those located at least 1.5 µm away from the nucleus toward the culture medium side were designated as ‘Floating’, those present on the cell adhesion surface were designated as ‘Close to glass’, and those adhering to the nucleus were designated as ‘Close to cells’.

### Statistical analysis

Data are presented as mean ± standard error of the mean (SEM). Statistical significance between two groups was assessed using a two-tailed unpaired Student’s *t*-test. For comparisons among multiple groups, one-way analysis of variance followed by appropriate post hoc tests were performed. Differences were considered statistically significant at *p* values of < 0.05. All statistical analyses were conducted using GraphPad Prism software (GraphPad Software, Inc., San Diego, CA, USA).

## Results

### PANC-1 clones derived from solitary culture exhibited enhanced ciliogenesis and distinct ciliary morphologies

We utilized two previously established PANC-1 clones (clone #70 and #76) (Fig. S1A), which were characterized in our prior study as exhibiting enhanced cell mass formation in a ciliogenesis-dependent manner (Shirakawa *et al*., 2025). To investigate whether primary cilia-derived EVs contribute to this phenotype, we first optimized culture conditions to minimize interference from serum-derived factors. In our previous study, PANC-1 cells were cultured in medium containing 10% FBS (Shirakawa *et al*., 2025). Here, we examined whether clone #70 and #76 could maintain their enhanced ciliogenesis capacity under serum-deprived conditions. Cells were grown to confluency and then cultured for an additional 24 h in medium containing 1% EV-depleted FBS (pre-cleared by ultracentrifugation), after which most subsequent analyses were performed (Fig. S1B). Under this condition, both clone #70 and clone #76 displayed a significantly higher proportion of ciliated cells compared with parental PANC-1 cells: 42.0 ± 1.7% and 44.2 ± 3.9%, respectively, versus 29.3 ± 2.0% in the parental line (Fig. S1C, S1D). Based on these results, we adopted this low-serum (1% EV-depleted FBS) culture condition to induce ciliogensis in all following experiments. Primary cilia and their basal bodies were reliably visualized by staining for ARL13B and FOP (also known as CEP43), respectively (Fig. S1E).

We also examined the positions of primary cilia formed in clone #70. Primary cilia were classified into three types: **‘Top’** (cilia protruding from the medium-side cell surface), **‘Bottom’** (cilia located between the cell and the basement glass surface), and ‘**Middle**’ (cilia protruding between adjacent cells) (Fig. S1F). Nearly half of the primary cilia protruded from the medium-side cell surface, while approximately 20% were located between the cell and the glass surface (Fig. S1G; Top: 46.8 ± 0.7%; Bottom: 19.2 ± 1.5%). The remaining ∼30% of primary cilia projected between cells (Fig. S1G; Middle: 34.0 ± 2.0%).

High-magnification imaging revealed numerous ARL13B-positive particles in clone #70 (Fig. 1A, arrowheads). In addition, cilia with a bulged morphology at their tips were frequently observed (Fig. 1A, arrow in enlarged image), as well as branched cilia (Fig. 1A, arrow in enlarged image), suggesting that PANC-1 clones surviving solitary culture conditions acquire diverse ciliary structural characteristics associated with EV emission. We thus performed a detailed morphological analysis of primary cilia, categorizing them into five distinct types: (1) ‘**Normal’**, (2) ‘**Bulged’**, characterized by a swollen ciliary tip, (3) ‘**Emission’**, defined by the presence of particles in close proximity to the cilium, (4) ‘**Very short’**, consisting of markedly shortened cilia (including those oriented perpendicular to focal plane), and (5) ‘**Branched’**, featuring two cilia protruding from a single basal body (Fig. 1B). The ‘Normal’ type predominated in all cell lines; however, its proportion was slightly but significantly reduced in clone #70 and #76 (Parental: 86.0 ± 1.0% vs. clone #70: 70.4 ± 3.8%, *p* = 0.012; vs. clone #76: 72.2 ± 3.6%, *p* = 0.012)(Fig. 1C). In contrast, the ‘Bulged’ type accounted for more than 15% of cilia in clones #70 and #76, significantly higher than in parental PANC-1 cells (Parental: 6.8 ± 1.3% vs. clone #70: 15.7 ± 2.4%, *p* = 0.020; vs. clone #76: 16.1 ± 2.2, *p* = 0.020)(Fig. 1C). Although less abundant overall, the ‘Emission’ type was also significantly enriched in clones #70 and #76 (Parental: 1.3 ± 0.2% vs. clone #70: 3.4 ± 0.6%, *p* = 0.062; vs. clone #76: 4.0 ± 1.1%, *p* = 0.044)(Fig. 1C). The ‘Very short’ type was slightly more frequently in clone #70 and #76 (Parental: 5.2 ± 1.3% vs. clone #70: 10.0 ± 1.8%, *p* = 0.098; vs. clone #76: 7.4 ± 1.4, *p* = 0.323)(Fig. 1C). The ‘Branched’ type was extremely rare (<1%) in both parental PANC1, clone #70 and #76 (Fig. 1C).

**Figure 1.**
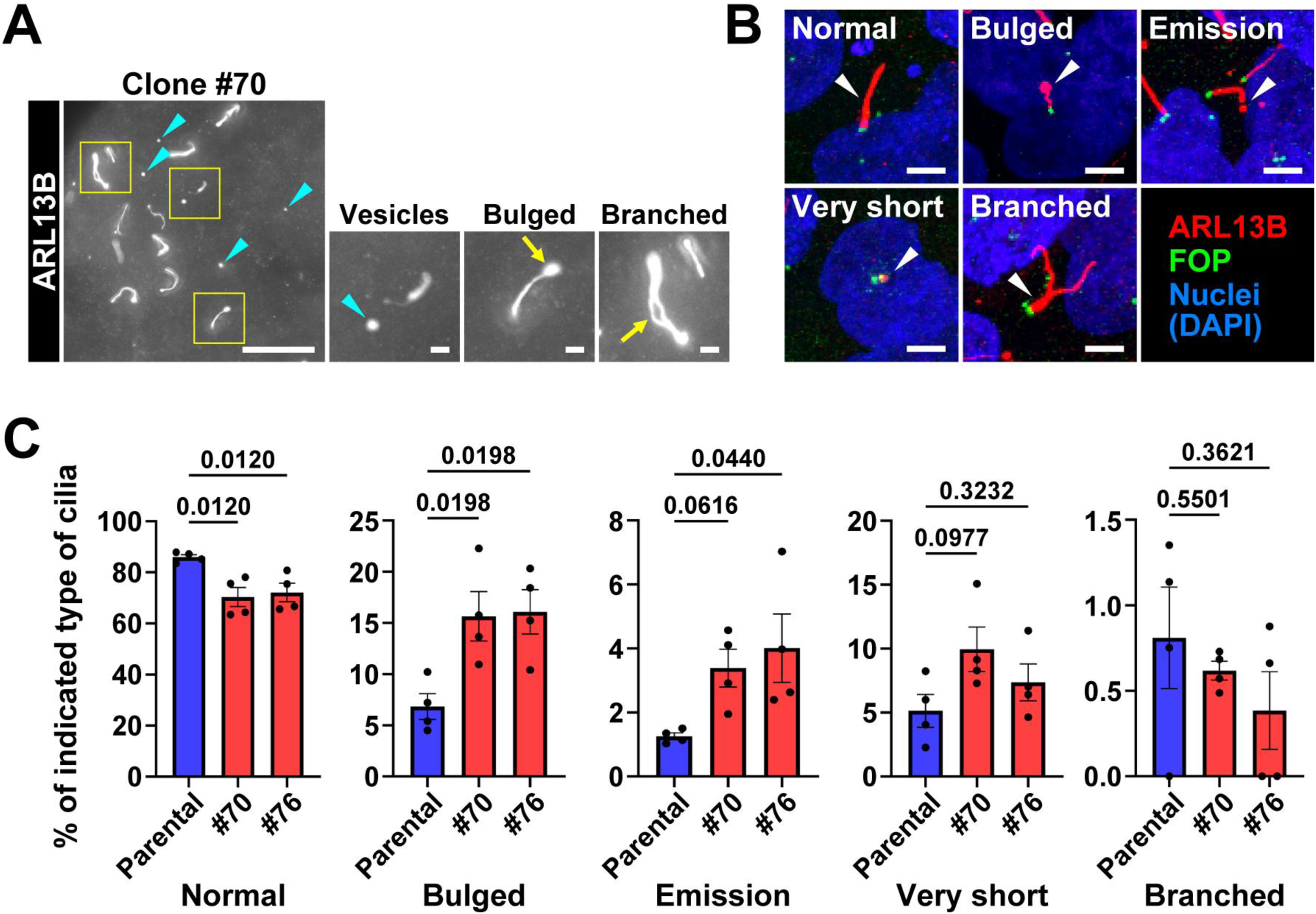
Enhanced EV-related structures and morphological diversity of primary cilia in PANC-1 clones derived from solitary culture. **(A)** Representative immunofluorescence images showing branched cilia (arrow), bulged ciliary tips (arrow), and vesicles (arrowheads) in clone #70 cells. Scale bar, 20 µm (main image) and 2 µm (enlarged views). **(B)** Representative immunofluorescence images showing classification of five primary cilium morphologies (arrowheads). Primary cilia are detected with ARL13B (red), centrioles with FOP (green), and nuclei with DAPI (blue). Scale bars, 5 µm. **(C)** Quantification of each ciliary morphology. Data represent mean ± SEM from five fields of view in each of four independent experiments. Statistical significance was defined as *p* < 0.05, determined using one-way ANOVA.

To further characterize the ‘Bulged’ type, we performed double immunofluorescence staining for ARL13B and acetylated-α-tubulin (Ac-tub), a major component of the ciliary axoneme. Based on Ac-tub staining, ‘Bulged’-type cilia were divided into three subtypes: **‘Ac-tub-negative’** (absence of Ac-tub signal within the bulge), **‘Ac-tub-positive’** (Ac-tub signal extending into the bulge), and **‘Circle’** (forming a ring-like tip structure)(Fig. S2A). The ‘Ac-tub-negative’ subtype was the most prevalent, and this trend was found to be statistically significant (Fig. S2B).

### Primary cilia-derived EVs, including a larger population of floating EVs, increase in PANC-1 clones derived from solitary culture

Given that PANC-1 clones derived from solitary culture exhibited more than twofold increase in primary cilia with bulged tips or ARL13B-positive particles compared with parental cells (Fig. 1C), we next quantified putative primary cilia-derived EVs in confocal fluorescence images. To exclude the possibility that the ARL13B-positive particles represented nonspecific signals, we performed double immunofluorescence staining using two independent primary antibodies: a rabbit polyclonal antibody (17711-1-Ap) and a mouse monoclonal antibody (66739-1-Ig). All primary and secondary antibodies were centrifuged prior to use to eliminate aggregates and false-positive puncta. ARL13B-positive particles were consistently labeled by both antibodies, regardless of their distance from the primary cilium, confirming the specificity of the signals (Fig. 2A).

**Figure 2.**
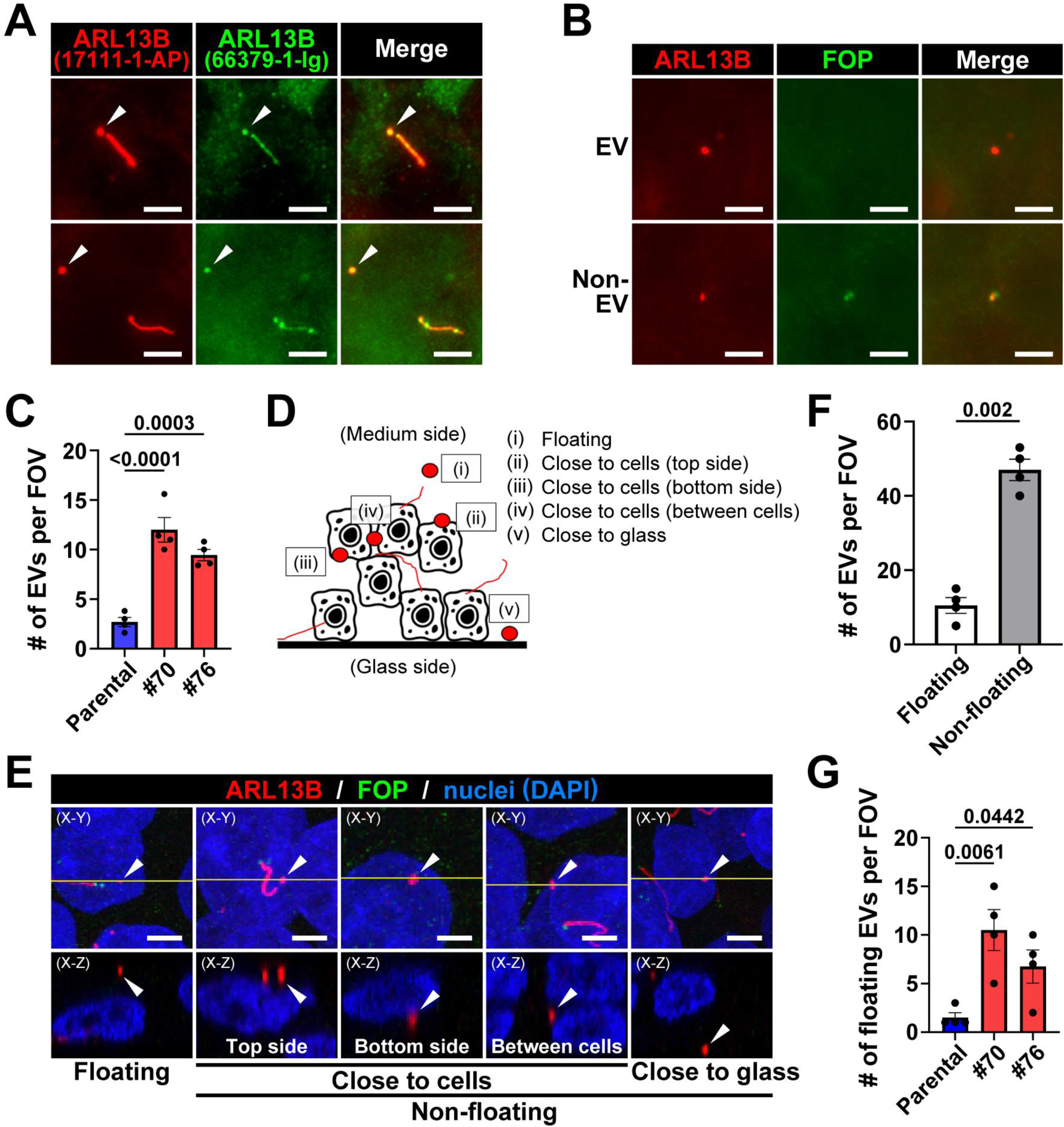
Increased production and spatial dispersion of cilia-derived EVs by PANC-1 clones derived from solitary culture. **(A)** Immunostaining of vesicles and primary cilia using anti-ARL13B antibodies (red: 17711-1-AP; green: 66379-1-Ig). Scale bar, 5 µm. **(B)** Immunostaining for ARL13B (red) and FOP (green) to define EVs. EVs were defined as ARL13B-positive particles located away from FOP staining, whereas particles merged with or adjacent to FOP were classified as non-EVs. Scale bar, 5 µm. **(C)** Quantification of EV numbers in parental PANC-1 cells, clone #70 and #76. Data represent mean ± SEM from five fields of view (FOV) in each of four independent experiments. **(D)** Schematic illustration of the classification of EV locations. **(E)** Representative immunofluorescence images showing the five EV spatial categories. Upper: maximum-projection x-y images. Lower: x-z cross-sectional images taken at the yellow line in x-y images. Scale bar, 20 µm. **(F)** Quantification of ‘floating EVs’ and ‘non-floating EVs’ in clone #70. Data represent mean ± SEM from five fields of view (FOV) in each of four independent experiments. **(G)** Quantification of floating EVs in parental PANC-1 cells, clone #70 and #76. Data represent mean ± SEM from five fields of view (FOV) in each of four independent experiments. Statistical significance was defined as *p* < 0.05 and assessed using one-way ANOVA (panels C and G) or two-tailed unpaired Student’s *t*-test (panel F).

Since very short or truncated primary cilia can appear similar to ARL13B-positve puncta, we established a conservative criterion for EV identification. ARL13B-positive particles located in close proximity to FOP, a marker of the basal body, were classified as short or truncated cilia and excluded from quantification (Fig. 2B). Under this definition, ARL13B-positive particles positioned away from FOP were counted as primary cilia-derived EVs in randomly selected fields of view. Quantitative analysis revealed that the number of primary cilia-derived EVs was significantly higher in both clone #70 and clone #76 compared with parental PANC-1 cells (Parental: 2.7 ± 0.5 vs. clone #70: 12.0 ± 1.1, *p* < 0.001; vs. clone #76: 9.5 ± 1.0, *p* < 0.001)(Fig. 2C). These results indicate that the primary cilia in solitary condition-surviving PANC-1 clones possess a markedly enhanced capacity for EV release.

To further characterize where primary cilia-derived EVs reside within the cell culture environment, we analyzed their spatial distribution in confocal images. EVs were categorized into three groups: ‘**floating’**, ‘**close to cells’** (including above, beneath, or between cells), and ‘**close to glass’** (Fig. 2D, 2E). In clone #70, which exhibited the highest overall ARL13B-positive EV number (Fig. 2C), the number of ARL13B-positive EVs classified as ‘non-floating’ (i.e. close to cells or close to glass) was significantly greater than the number of floating EVs (floating: 10.5 ± 2.1 vs. non-floating: 47.0 ± 2/9; *p* =0.002)(Fig. 2F). The proportion of floating EVs was also significantly higher in clone #70 and #76 than in parental PANC-1 cells (Parental: 1.5 ± 0.5 vs. clone #70: 10.5 ± 2.1, *p* = 0.006; vs. clone#76: 6.8 ± 1.7, *p* = 0.04)(Fig. 2G). These findings suggest that PANC-1 cells that survived solitary culture conditions release a greater number of EVs capable of dispersing freely into the culture medium compared with parental PANC-1 cells.

### PANC-1 clones derived from solitary culture selectively increase the release of primary cilia-derived EVs

Given that PANC-1 clones derived from solitary culture exhibited an increased number of putative freely dispersing ARL13B-positive EVs, we hypothesized that ARL13B-positive EVs would be recovered more abundantly from PANC-1 clones derived from solitary condition cultures than from parental PANC-1 cell cultures. To quantify primary cilia-derived EVs release, we collected EVs from conditioned media using ultracentrifugation. Parental PANC-1 cells, clone #70 and #76 were grown to confluency and then cultured for an additional 24 h in medium containing 1% pre-cleared FBS prior to media collection. After sequential centrifugation steps to remove dead cells and debris, EV-containing pellets were obtained by ultracentrifugation (Fig. 3A). Both the EV pellets and corresponding cell layers remaining after media removal were solubilized in SDS-PAGE sample buffer and analyzed by Western blotting for ARL13B (a primary cilia marker), ALIX (an exosome marker), and GAPDH (a cytoplasmic protein)(Fig. 3B). The release level of each protein was evaluated by calculating the ratio of its abundance in the EV pellet to that in the corresponding cell lysate (Fig. 3C). GAPDH was barely detectable in the EV pellets, indicating minimal contamination from cytoplasmic proteins and confirming successful EV isolation. The ratios of GAPDH in EV pellets to cell lysates were similarly low (∼0.04) across all groups (Parental: 0.037 ± 0.005 vs. clone #70: 0.044 ± 0.006, *p* = 0.836; vs. clone #76: 0.041 ± 0.010%, *p* = 0.836)(Fig. 3B, 3C). Notably, ARL13B levels in the EV pellets were significantly higher in clones #70 and #76 than in parental PANC-1 cells (Parental: 0.55 ± 0.03 vs. clone #70: 1.19 ± 0.12, *p* = 0.001; vs. clone #76: 1.09 ± 0.08, *p* = 0.002)(Fig. 3B, 3C). In contrast, ALIX levels in EV pellets did not differ significantly among the three cell types (Parental: 1.49 ± 0.18 vs. clone #70: 1.59 ± 0.15, *p* = 0.649; vs. clone #76: 1.78 ± 0.11, *p* = 0.353)(Fig. 3B, 3C). Together, these results demonstrate that PANC-1 cells surviving solitary culture conditions selectively enhance the release of primary cilia-derived EVs, rather than increasing EV production globally.

**Figure 3.**
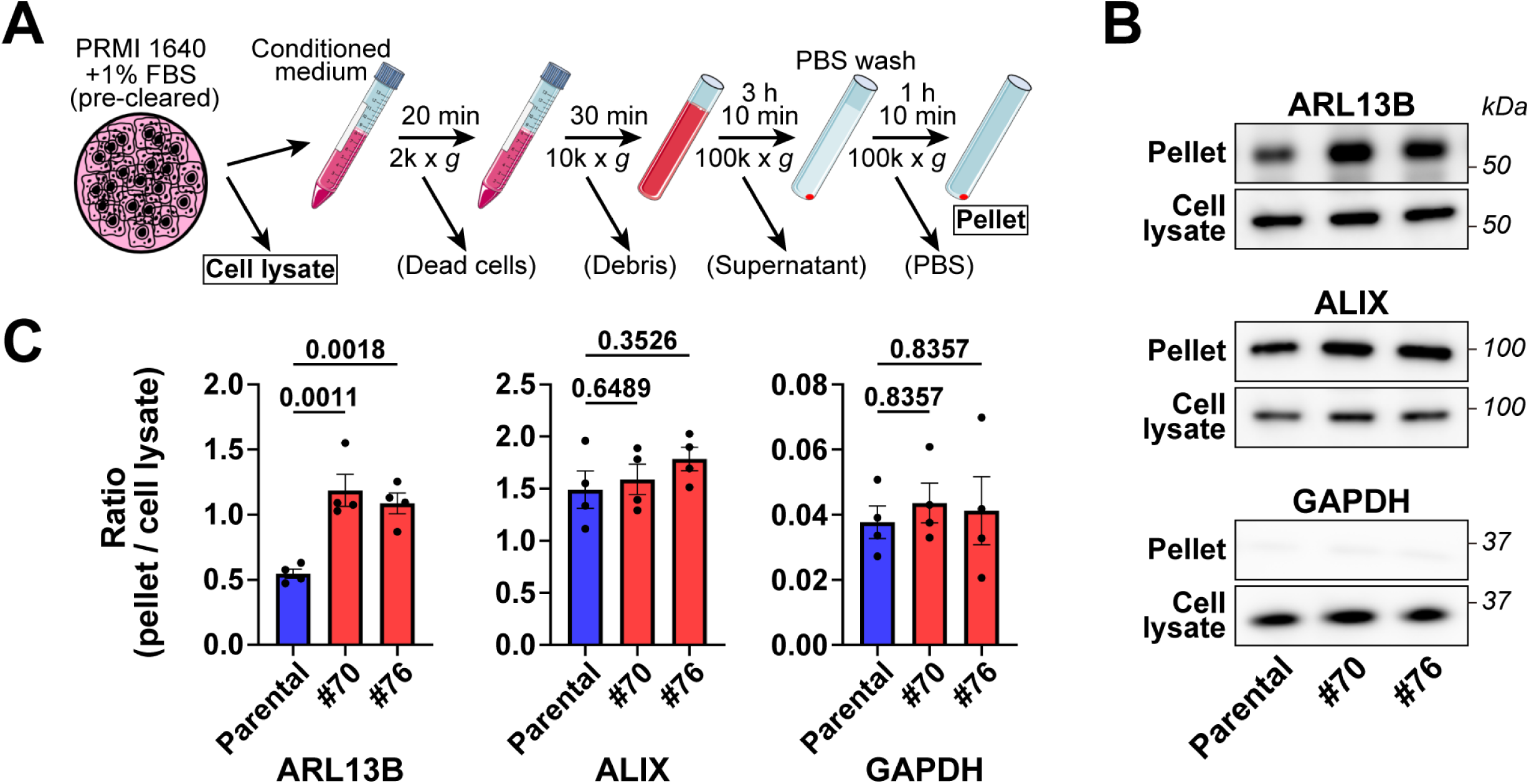
Selective enhancement of primary cilia-derived EV release in PANC-1 clones derived from solitary culture. **(A)** Schematic overview of the procedure used for EV collection. **(B)** Representative Western blots showing ARL13B, ALIX, and GAPDH levels in cell lysates and in EV pellets obtained from the conditioned medium of parental PANC-1 cells, clone #70 and #76. **(C)** Quantitative analyses of the pellet-to-cell lysate ratio for each protein (ARL13B, ALIX, and GAPDH). Data represent mean ± SEM from four independent experiments. Statistical significance was defined as *p* < 0.05 and assessed using one-way ANOVA.

### EVs enriched in primary cilia-derived EVs suppress cell migration and promote cell aggregation

We next sought to determine how the increased abundance of primary cilia-derived EVs in PANC-1 clones derived from solitary cultures affects cellular behaviors. However, both knockdown and knockout approaches proved unsuitable for functional disruption of ciliogenesis in our system. In knockdown experiments (Fig. S3A-S3C), siRNA-mediated suppression of *IFT88* or *KIF3A* expression unexpectedly did increase the amounts of GAPDH, ARL13B, and ALIX detected in EV pellets compared with siNC-treated controls (Fig. S3D). These results suggest that inhibiting ciliogenesis triggers unintended leakage of cytoplasmic components into the culture medium, complicating the interpretation of EV-related phenotypes. Furthermore, in CRISPR/Cas9-mediated knockout experiments, we were unable to isolate any viable clones deficient in either *IFT88* or *KIF3A*, despite screening more than 50 clones for each gene. These repeated failures prevented us from performing gene disruption-based analyses of ARL13B-positive EVs.

Given these limitations, we instead compared the biological effects of EVs collected from clones #70 and #76 with those from parental PANC-1 cell cultures (Fig. 4A). To assess the impact of EVs on cell migration, we performed wound-healing assays. Confluent parental PANC-1 monolayers were scratched, and the medium was replaced with medium in which EVs collected from either the parental PANC-1 cells, clone #70 or #76 cultures had been resuspended (Fig. 4A). The recovered area was measured 48 h after wounding (Fig. 4B). Parental PANC-1 cells treated with EVs derived from clone #70 and #76 exhibited significantly reduced wound closure compared with those treated with parental EVs (Parental EVs: 9.9 ± 1.0% vs. clone #70 EVs: 7.5 ± 0.8%, *p* = 0.086; vs. clone #76 EVs: 5.5 ± 0.9%, *p* = 0.015)(Fig. 4B, 4C). These results indicate that primary cilia-derived EVs suppress PANC-1 cell migration.

**Figure 4.**
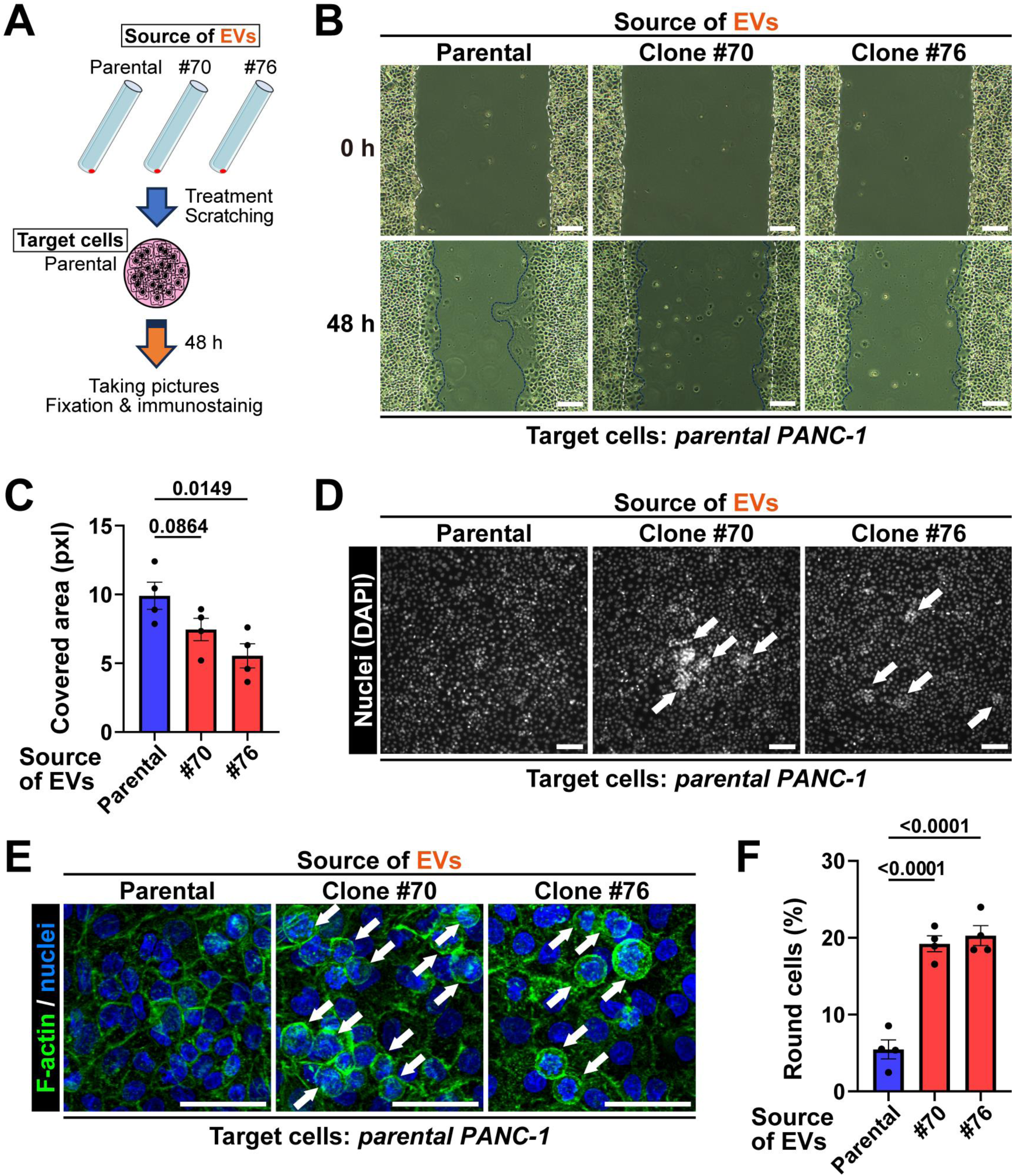
Suppression of cell migration and promotion of cell aggregation by PANC-1 clones-derived EVs. **(A)** Schematic overview of the experimental procedure for treating parental PANC-1 cells with primary cilia-derived EV-enriched EVs. **(B)** Representative phase-contrast images of wound closure in parental PANC-1 cells treated with EVs collected from parental PANC-1 cells, clone #70 or #76. Images were captured at 0 h and 48 h after wound creation. The initial wound edge (0 h) is indicated by white lines, and the boundary of the recovered area at 48 h is indicated by blue lines. Scale bars, 200 µm. **(C)** Quantification of recovered wound area at 48 h in parental PANC-1 cells treated with EVs from parental PANC-1 (blue) or clone #70 or #76 (red), presented in pixels (pxl). Data represent mean ± SEM from four independent experiments. **(D)** Fluorescence images of nuclei (DAPI) in parental PANC-1 cells treated with EVs from parental PANC-1 cells, clone #70 or #76 cultures for 48 h. Arrows indicate aggregated cells. Scale bars, 200 µm. **(E)** Representative fluorescence images showing cell morphology in parental PANC-1 cells treated with EVs from parental PANC-1 cells, clone #70 or #76 cultures for 48 h. Arrows indicate round-shaped cells. Scale bars, 50 µm. **(F)** Quantification of round-shaped cells in parental PANC-1 cultures treated with EVs from parental PANC-1 (blue) or clone #70 or #76 (red) for 48 h. Data represent mean ± SEM from five fields of view in each of four independent experiments. Statistical significance was defined as *p* < 0.05 was defined as and assessed using one-way ANOVA.

We next examined the effects of EVs on cell aggregation, given our previous observation that clones #70 and #76 exhibit enhanced cell mass formation (Shirakawa *et al*., 2025). Parental PANC-1 cells were treated with EVs derived from parental PANC-1 cells, clone #70 or #76 cultures, and nuclei were visualized by DAPI staining 48 h after treatment. Notably, cells treated with PANC-1 clones EVs formed more compact aggregates than those treated with parental-cell EVs (Fig. 4D). To further characterize the morphology of aggregated cells, we stained F-actin using fluorophore-conjugated phalloidin (Fig. 4E). The proportion of rounded PANC-1 cells was significantly higher in cultures treated with clone #70 and #76 EVs compared with parental PANC-1 EVs (Parental PANC-1 EVs: 5.5 ± 1.2% vs. clone #70 EVs: 19.2 ± 1.0%, *p* < 0.001; vs. clone #76: 20.3 ± 1.3%, *p* < 0.001)(Fig. 4F). Together, these results demonstrate that primary cilia-derived EVs from PANC-1 clones derived from solitary cultures not only inhibit migration but also promote cell aggregation and induce rounded cell morphology, implicating them as functional mediators of cell behavioral changes associated with solitary condition-surviving PANC-1 clones.

## Discussion

One of the most important findings of this study is that PANC-1 clones, which were grown from a single cell and exhibited an enhanced capacity of primary cilia formation, showed increased release of primary cilia-derived EVs. In the PANC-1 clones derived from solitary condition culture, we observed numerous ARL13B-positive particles by immunofluorescence microscopy (Fig. 1 and 2). Since the identification of ARL13B-positive puncta could be subject to alternative interpretations, we implemented three rigorous experimental procedures to validate our conclusion. First, to eliminate the possibility that immunoglobulin aggregates or nonspecific fluorescent particles from the primary and secondary antibodies contributed to the observed signals, we centrifuged all antibodies at 20,000 **×** g for 30 min prior to use. Second, we performed dual immunostaining with two independent anti-ARL13B antibodies, ensuring that only particles labeled by both antibodies were considered ARL13B-positive. Third, to exclude the possibility that extremely short primary cilia might be misidentified as ARL13B-positive EVs, we classified ARL13B-positive particles as EVs only when they were spatially separated from the centriole. Together, these carefully designed controls strongly support our interpretation that the ARL13B-positive particles observed in the immunostaining analyses present primary cilia-derived EVs.

A high frequency of primary cilia with bulged tips was also observed in PANC-1 clones derived from solitary culture (Fig. 1). Primary cilia-derived EVs have been reported to be released through the accumulation of vesicular contents at the ciliary tip followed by pinching off, a process that depends on actin polymerization (Phua *et al*., 2017). Our data indicate that the increase in primary cilia-derived EVs in PANC-1 clones culture cannot be attributed solely to the higher number of primary cilia; rather, the EV-release capacity of these cells also appears to be enhanced. Specifically, the number of ARL13B-positive particles increased by approximately threefold in PANC-1 clones compared with parental PANC-1 cells (Fig 2C), a difference that corresponds well to the combined effect of the ∼1.5-fold increase in ciliogensis (Fig. S1D) and the ∼2-fold increase in buldged-tip frequency (Fig. 1C). We further demonstrated that most ciliary tip bulges in clone #70 lacked acetylated tubulin (Fig. S2). This observation is consistent with our previous findings and other reports showing that EVs derived from primary cilia do not contain acetylated tubulin, a major structural component of the ciliary axoneme (Nager *et al*., 2017; Phua *et al*., 2017). Taken together, these results strongly support the interpretation that the bulges observed at ciliary tips in PANC-1 clones represent structures that are released as primary cilia-derived EVs.

Interestingly, although rare, we observed acetylated tubulin forming a ring-like (‘Circle’) structure at the distal ends of some primary cilia in clone #70 (Fig. S2). Such an unusual morphology has not been previously reported and might reflect aberrant structural features associated with the cancer cell state. In addition, we identified primary cilia that were split into two branches (‘Branched’; Fig. 1). This phenomenon might not be specific to a single cell type, as human pancreatic islet cells have been shown by electron microscopy to possess multiple ciliary projections emerging from a single basal body (Polino *et al*., 2023). Similarly, RPTEC/TERT1, hTERT-immortalized renal proximal tubule epithelial cells, have been reported to occasionally exhibit primary cilia extending in two directions from a single centriole, resembling the branched form (Hansen *et al*., 2025). In contrast, to our knowledge, the presence of such branched primary cilia in pancreatic cancer tissue has not been reported previously. These atypical morphological features, including branched cilia and circular ciliary tips, were extremely rare, being observed in less than 1% of primary cilia in PANC-1 cells. Further studies are required to determine their biological significance and potential relevance in oncological and pathological contexts.

A greater number of primary cilia-derived EVs were detected on or between cells in clone #70 (Fig. 2F). This observation suggests that EVs released from PANC-1 clones derived from solitary culture increased cell surface retention and readily associate with the cell membrane. Consistent with this interpretation, immunostaining revealed a markedly larger increase in EV-associated ARL13B-positive particles (3-4-fold higher in PANC-1 clones than in parental PANC-1 cells; Fig. 2) than that observed by biochemical analysis of EV pellets from clone culture media, which showed an approximately twofold increase in ARL13B levels (Fig. 3). These findings support the notion that a substantial fraction of primary cilia-derived EVs remains attached to the cell surface of PANC-1 clones. This interpretation is further supported by our results that, in clone #70, less than half of the primary cilia extended their tips toward the culture medium, while the majority projected either into intercellular spaces or toward the bottom glass surface (Fig. S2). Such spatial orientation of primary cilia may contribute to the preferential accumulation of primary cilia-derived EVs between cells in clone #70. Nonetheless, we cannot exclude the possibility that primary cilia-derived EVs initially released into intercellular spaces or at the cell-glass interface subsequently migrate or are displaced toward medium-facing cell surface as a result of dynamic movements of stacking cells.

An alternate explanation, however, is that PANC-1 clones themselves possess increased surface adhesiveness. Notably, elevated adhesiveness of EVs or of the cell surface is a known feature of mesenchymal cell types. For example, ARL13B-positive cilia-derived EVs from fibroblasts such as NIH/3T3 cells or mouse embryonic fibroblast, both exhibiting mesenchymal characteristics, frequently remain attached to the plasma membrane (Phua *et al*., 2017). Our previous work identified mesenchyme-like traits in PANC-1 clones that have survived solitary culture conditions, suggesting that increased cellular adhesiveness might be an intrinsic property of these clones. The enhanced adhesion of primary cilia-derived EVs released from PANC-1 clones might have implication for the early metastatic behavior of PDAC. Cell adhesion has been recognized as a critical determinant of tumor metastasis, including the adhesive properties of EV surfaces and associated adhesion molecules (Jerabkova-Roda *et al*., 2022). These findings raise the possibility that the strong adherence of cilia-derived EVs contributes to the mechanisms underlying the remarkably early metastasis characteristics of PDAC, one of the malignancies with the poorest prognosis (Kuehn, 2020; Sung *et al*., 2021).

We collected EVs released into the culture medium by ultracentrifugation and analyzed their components by Western blotting. The results showed that the amount of ARL13B was significantly increased in the EV pellets derived from PANC-1 clones derived from solitary cultures compared with those from parental PANC-1 cells (Fig. 3). These findings provide strong evidence that the release of primary cilia-derived EVs is elevated in PANC-1 clones. In contrast, the amount of ALIX (apoptosis-linked gene 2-interacting protein X) present in EV pellets was unchanged between PANC-1 clones and parental PANC-1 cultures (Fig. 3). ALIX is a component of the endosomal sorting complex required for transport (ESCRT) complex and is widely used as a marker for exosomes (Dowlatshahi *et al*., 2012; Matsuo *et al*., 2004). The absence of changes in ALIX levels therefore suggests that the increase observed in PANC-1 clones is specific to primary cilia-derived EVs rather than a global increase in EV production. Further investigation will be required to elucidate the mechanism underlying this selective enhancement of cilia-derived EV release.

We were unable to apply knockdown approaches effectively in the present study, despite the success in suppressing primary cilia formation in PANC-1 cells by siRNA-mediated knockdown of *KIF3A* or *IFT88* (Fig. S3). In clone #70, knockdown of either *KIF3A* or *IFT88* did not reduce the release of primary cilia-derived EVs into the culture medium (Fig. S3). Instead, the amounts of ARL13B, ALIX, and GAPDH detected in the culture medium were all increased compared with control siRNA-treated cells. These results suggest that inhibition of ciliogenesis in PANC-1 clones derived from solitary culture elicits unexpected cellular responses that mask or counteract the intended suppression of cilia-derived EV release. An alternative explanation is that primary cilia-derived EVs were not efficiently recovered from siRNA-treated cultures using the three-step centrifugation protocol employed in this study. One plausible scenario is that the lipofection reagent used for siRNA delivery interfered with recovery of primary cilia-derived EVs, as lipofection reagents have been reported to perturb the detection and analysis of EV components (Roerig and Schulz-Siegmund, 2023). Given their strong membrane-adhesive properties, lipofection reagents might increase nonspecific interactions between EVs and cellular or plastic surfaces, leading to loss of putatively ‘sticky’ primary cilia-derived EVs during centrifugation. EVs isolation protocols will therefore be required to reliably collect primary cilia-derived EVs from cultures subjected to lipofection-based treatments.

The negative impact of disrupting key ciliogenic genes was even more evident in our attempts at gene knockout. We were unable to obtain *KIF3A*- or *IFT88*-knockout PANC-1 cells using CRISPR/Cas9, despite repeated attempts. This difficulty is consistent with several previous reports in which targeted gene knockout in PANC-1 cells was also failed, even when using CRISPR/Cas9 systems (Lentsch *et al*., 2019; Meng *et al*., 2025; Ramaker *et al*., 2021). Other groups have similarly relied on knockdown rather than knockout strategies when investigating primary cilia-related genes in PANC-1 cells (Kobayashi *et al*., 2017; Mashima *et al*., 2022). To our knowledge, no studies have successfully suppressed primary ciliogenesis in PANC-1 cells by knocking out cilia-related genes. These collective observations suggest that primary cilia-related genes are essential for PANC-1 cell viability and that complete loss of these genes likely causes severe cellular damage or lethality, thereby preventing the establishment of viable knockout clones.

Several previous studies have investigated the effects of EVs, including exosomes, on cell migration. Multiple reports indicate that EVs function as chemoattractants in cancer cells, thereby promoting cell migration (Sung *et al*., 2015; Sung and Weaver, 2017). In contrast, other studies have demonstrated that EVs can inhibit cell migration (Liu *et al*., 2023; Wu *et al*., 2017). Notably, EVs released from irradiated pancreatic cancer cells have been reported to suppress the migration and invasion of pancreatic cancer cells (Nakaoka *et al*., 2025). These findings suggest that the effect of EVs on cell migration might depend on their cellular origin or situation. In this study, we showed that EVs containing an increased proportion of primary cilia-derived EVs, collected from clone #70 and #76 cell culture media, suppress the migration of parental PANC-1 cells in a wound-healing assay (Fig. 4A and 4B). Since the wound-healing assay does not generate a concentration gradient of added EVs, the observed effects likely reflect changes in cell motility rather than chemotactic responses to primary cilia-derived EVs.

EVs derived from clone #70 and #76 also promoted the formation of cell masses, and the actin cytoskeleton of the cells within these masses exhibited a circular morphology (Fig. 4D). EVs have previously been reported to influence actin polymerization in recipient cells (Kapustin *et al*., 2025; McAtee *et al*., 2025). Consistent with these reports, our findings therefore suggest that primary cilia-derived EVs released from PANC-1 clones derived from solitary culture might modulate actin polymerization and consequently inhibit cell migration in recipient cells. Notably, EVs released from the primary cilia of retinal photoreceptor cells have been shown to contain actin-related 2/3 (Arp2/3) complex, which initiates actin polymerization (Spencer *et al*., 2019). Although the ingredients of EVs released from the primary cilia of PANC-1 cells remain to be determined, we hypothesize that these EVs modulate cell migration and morphology by directly or indirectly regulating actin-related signaling pathways in recipient cells. Further studies will be required to test this hypothesis and to identify the relevant EV components.

We have reported that primary cilia in PANC-1 cells play an essential role in cell mass formation (Shirakawa *et al*., 2025). The present study demonstrates that the primary cilia-dependent cell mass formation in PANC-1 cells is mediated, at least in part, by EVs derived by primary cilia. Primary cilia have been reported to harbor molecules involved in cell adhesion, including integrins (Praetorius *et al*., 2004; Goodman and Zallocchi, 2017). Based on these reports, we propose that primary cilia-derived EVs released from PANC-1 clones may carry such cell adhesion-related molecules and thereby promote cell mass formation. Although EV-mediated cell mass formation might also involve enhanced cell proliferation, the increased cell adhesion of cilia-derived EVs observed in Figure 2 is also likely to represent an additional contributing mechanism.

In conclusion, we demonstrated that pancreatic cancer cells derived from solitary conditions exhibit an increased release of primary cilia-derived EVs, and that this increase suppresses the migratory capacity of pancreatic cancer cells while promoting the formation of cell mass. Primary cilia in glioma cells have also been reported to release EVs, and although indirect effects cannot be excluded, these EVs enhanced glioma cell proliferation (Hoang-Minh *et al*., 2018). Despite such observations, very few studies have examined the functional roles of primary cilia-derived EVs, and none have explored their relevance in cancer biology with the exception of the glioma study. Our findings therefore represent one of the few reports elucidating the function and potential significance of primary cilia-derived EVs released from cancer cells. Further investigation will be required to clarify the mechanisms responsible for the increased production of cilia-derived EVs in PANC-1 cells surviving solitary conditions and to determine how these EVs regulate cell migration and cell mass formation.

## Acknowledgements

A part of this work was conducted in the Natural Science Center for Basic Research and Development at Hiroshima University (NBARD-00002) under the support from the MEXT Project to promote public utilization of advanced research infrastructure (Program for supporting the construction of core facilities; Grant Number JPMXS0441300023).

## Data Availability Statement

Data are available upon request from the corresponding author. Ethics approval was not required as this study involved only in vitro cell culture experiments.

## Author Contribution Statement

Conceptualization: TH, RN, KI

Funding acquisition: KI, KU

Investigation: TH, RN, KS, FI

Project administration: KU, ST, KI

Supervision: KI

Writing-original draft: TH, RN, KI

Writing-review & editing: all authors

## Figure Legends

**Figure S1.**
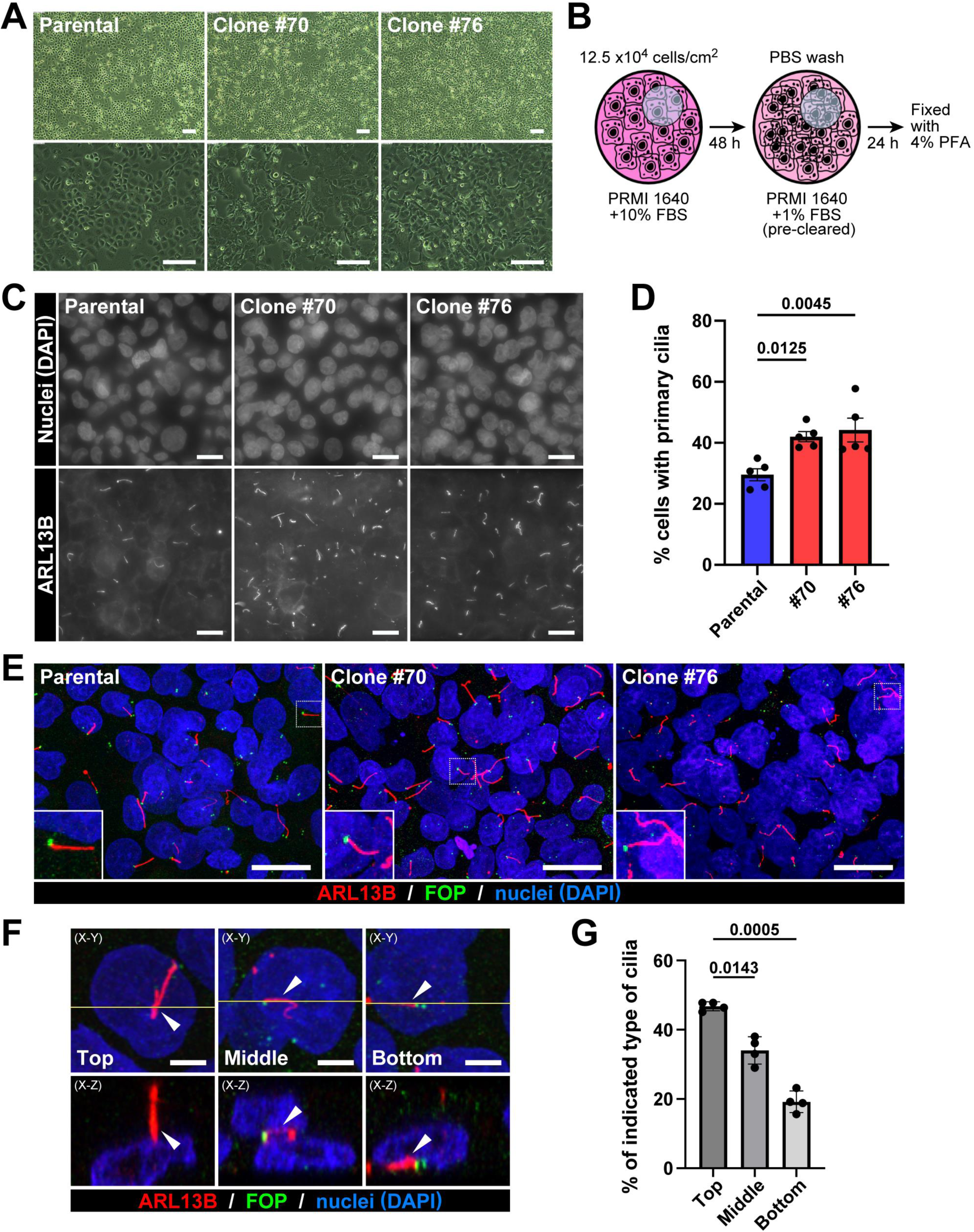
Validation of enhanced primary cilia formation in PANC-1 clones under low-serum conditions. **(A)** Representative phase-contrast images of parental PANC-1 cells, clone #70 and clone #76 acquired using an inverted microscope (DMI3000B, Leica) equipped with a 4× (top) and 10× (bottom) objective lenses. Scale bars, 200 µm. **(B)** Schematic overview of experimental conditions optimized to minimize nonspecific effects of serum-derived factors on experiments. **(C)** Representative immunofluorescence staining of nuclei (DAPI, upper) and primary cilia (ARL13B, lower) in parental PANC-1 cells, clone #70 and #76. Scale bars, 20 µm. **(D)** Quantification of the proportion of ciliated cells shown in (C). Data represent mean ± SEM from three independent experiments. Statistical significance was defined as *p* < 0.05 and assessed using one-way ANOVA. **(E)** Immunofluorescence staining of nuclei (DAPI, blue), primary cilia (ARL13B, red), and centrioles marking the basal boy (FOP, green) in parental PANC-1 cells, clone #70 or #76. Scale bars, 50 µm. **(F)** Representative immunofluorescence images showing three positions of primary cilia in clone #70. Primary cilia are labeled with ARL13B (red), centriole with FOP (green), and nuclei with DAPI (blue). Scale bars, 5 µm. **(G)** Quantitative analyses of cilia positions classified as shown in (F). Data represent mean ± SEM from five independent experiments.

**Figure S2.**
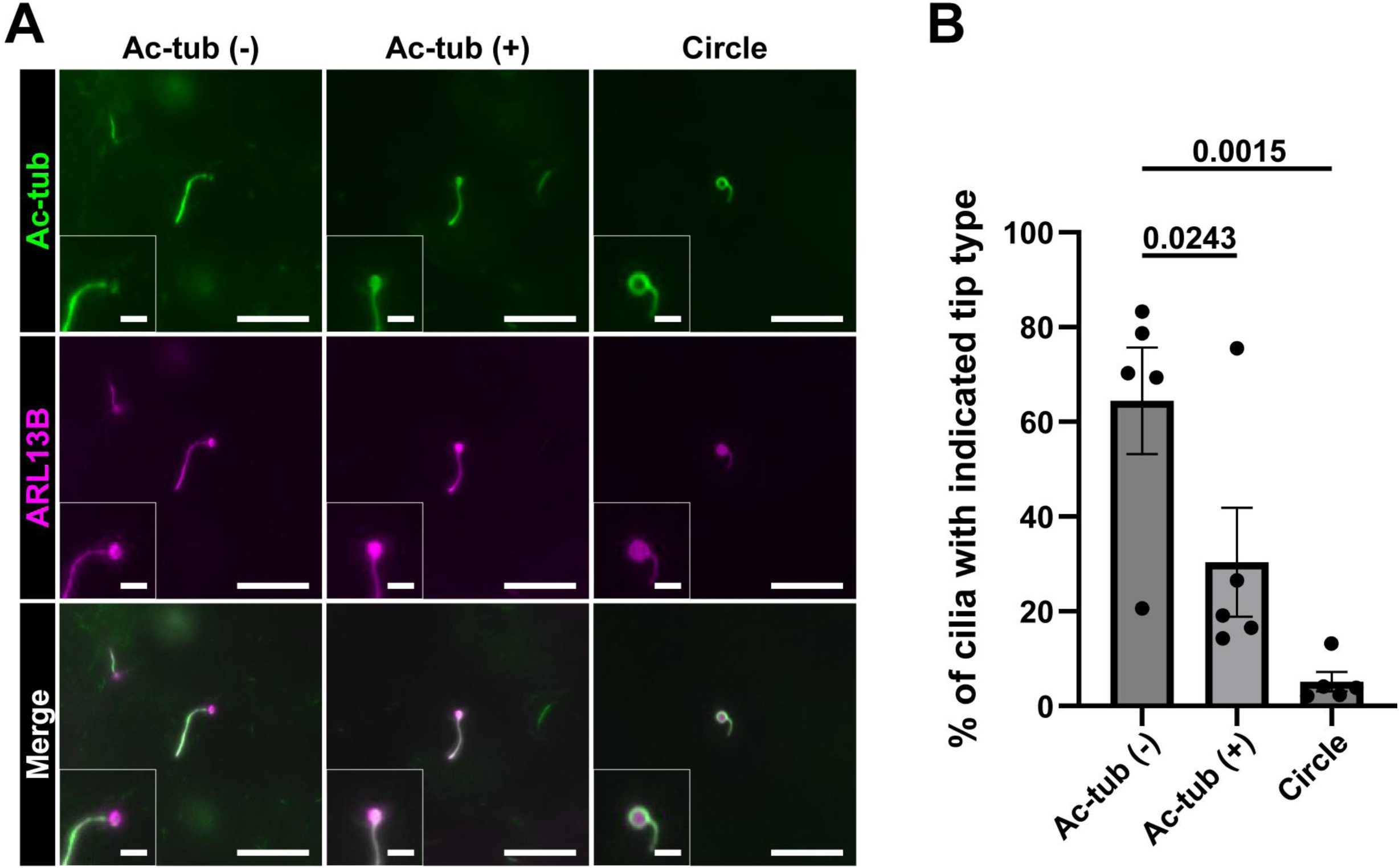
Three morphological types of bulged ciliary tips. **(A)** Representative immunofluorescence images showing three morphological types of bulged ciliary tips. Primary cilia are labeled with ARL13B (magenta), and ciliary axonemes with acetylated-α-tubulin (Ac-tub, green). Scale bars, 10 µm (main image) and 1 µm (insets). **(B)** Quantitative analysis of ciliary tip morphologies classified as shown in (A). Data represent mean ± SEM from five independent experiments. Statistical significance was defined as *p* < 0.05 and assessed using one-way ANOVA.

**Figure S3.**
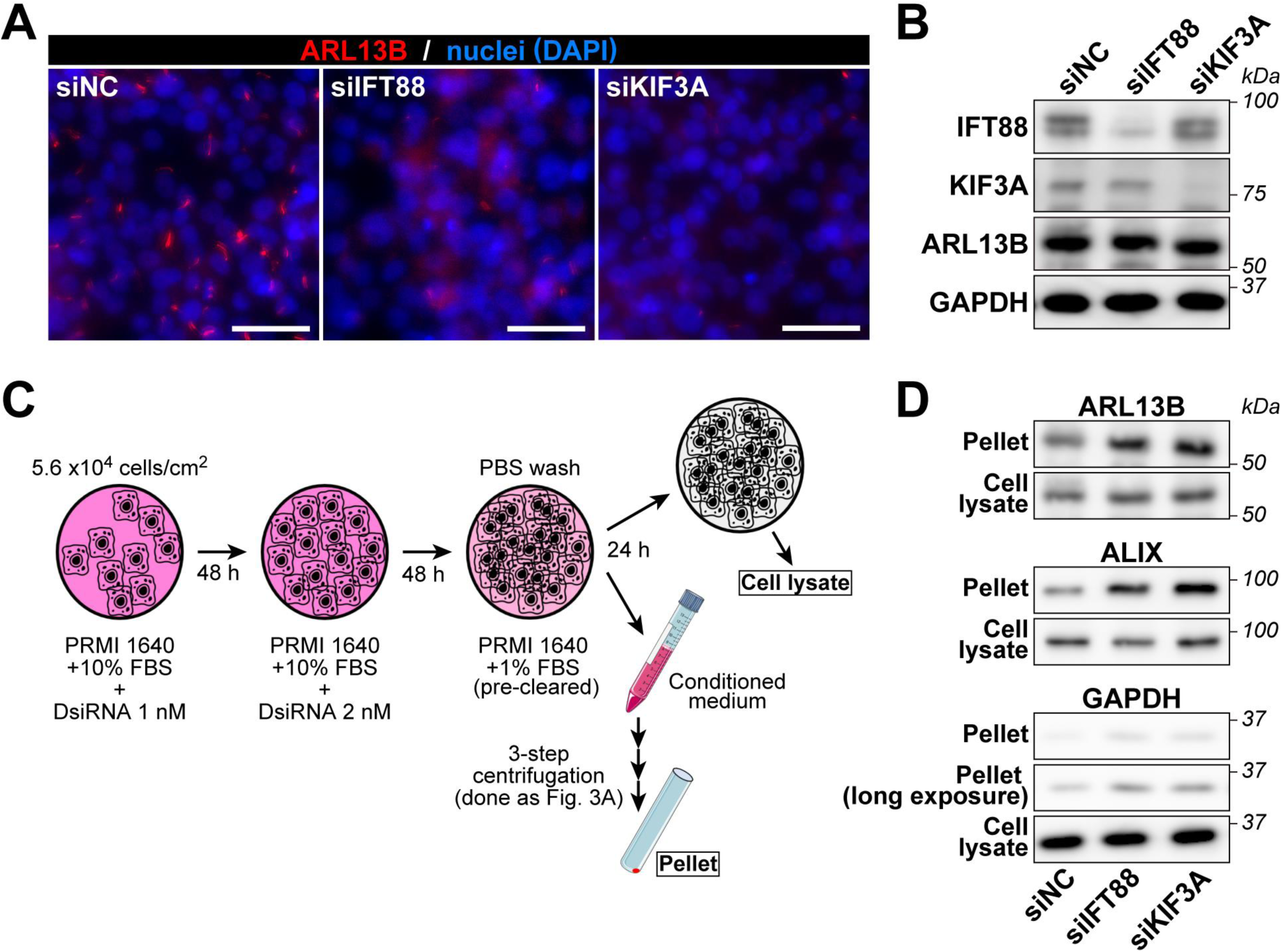
siRNA-mediated knockdown experiments. **(A)** Immunostaining for primary cilia (red, ARL13B) and nuclei (blue, DAPI) was performed in clone #70 following treatment with siRNAs targeting negative control, IFT88, or KIF3A. Immunofluorescence staining confirmed a marked reduction in the number of ciliated cells after IFT88 or KIF3A knockdown compared with negative control (siNC). Scale bars, 50 µm. **(B)** Western blots showing ARL13B, IFT88, KIF3A, and GAPDH protein levels in clone #70 after treatment with NC, IFT88, or KIF3A siRNAs. Western blot analysis also demonstrated a significant decrease in IFT88 and KIF3A protein expression by IFT88 and KIF3A knockdown, respectively, relative to siNC. **(C)** Schematic diagram of the experimental procedure for siRNA knockdown and subsequent EV collection. **(D)** Western blots of ARL13B, ALIX, and GAPDH in cell lysates and EV pellets collected from clone #70 cultures following treatment with NC, IFT88, or KIF3A siRNAs. Unexpectedly, siRNA-mediated suppression of IFT88 or KIF3A expression increased the amounts of GAPDH, ARL13B, and ALIX in EV pellets compared with siNC-treated controls.

